# A developmental chimera: co-option of appendage, secretory and myogenic programs underlies spider venom gland evolution

**DOI:** 10.64898/2026.07.10.737832

**Authors:** Giulia Zancolli, Afrah Hassan, Alistair P. McGregor, Yehu Moran, Marc Robinson-Rechavi

**Affiliations:** Department of Ecology and Evolution, University of Lausanne, 1015, Lausanne, Switzerland; Department of Biosciences, Durham University, South Road, Durham, DH1 3LE, United Kingdom; Department of Ecology, Evolution and Behavior, The Hebrew University of Jerusalem, 9190401 Jerusalem, Israel; Swiss Institute of Bioinformatics, 1015, Lausanne, Switzerland

## Abstract

The evolutionary emergence of complex organs requires the integration of distinct developmental modules, yet the mechanisms bridging divergent tissue programs into unified functional units remain poorly understood. Animal venom systems, combining high-output secretory epithelia with muscular delivery systems, a provide an exceptional model for studying this modular integration. Here, we show that the venom gland of the spider *Parasteatoda tepidariorum* is a developmental chimera built by the hierarchical co-option of three ancestral programs. The appendage-patterning gene *Distal-less* establishes the primary spatial coordinate system for gland development. Downstream, the transcription factor *sage* maintains secretory epithelium identity; its loss causes downregulation of key venom-processing enzymes and triggers a dramatic phenotypic reversion to a neuronal state. This neural upregulation, accompanied by the downregulation of Notch ligands (*delta* and *jagged*), reveals that *sage* actively represses a latent neural program to maintain secretory fate. Crucially, the myogenic factor *sum-1* governs the surrounding muscle layer to drive paracrine crosstalk (Wnt) and sustain lipid metabolism. Knockdown of *sum-1* downregulates *wntless* and fatty acid anabolism genes, alongside the abnormal accumulation of lipid droplets in the secretory epithelium. Ultimately, our findings demonstrate how a complex organ is assembled by fusing an appendicular address, a repressed neuroglandular program, and a myogenic metabolic driver. This modular architecture illustrates a fundamental principle in evolutionary developmental biology: novelty can arise by harnessing the physiological and metabolic capacities of adjacent tissues to fuel the physiological demands of a newly emerged organ.

**Significance statement:** How novel organs evolve by integrating distinct developmental modules remains a fundamental question in biology. Using the spider venom gland as a model, we show that this complex organ is a developmental chimera built by recruiting three ancestral tissue programs. The appendage-selector gene *Distal-less* (*Dll*) establishes the gland’s primary proximal-distal structural axis. Downstream, the transcription factor *sage* maintains secretory epithelial identity by actively repressing a default neural program. Concurrently, the myogenic regulator *sum-1* sustains epithelial lipid metabolism through paracrine Wnt signaling. These findings establish a mosaic model of organogenesis, demonstrating that radical evolutionary innovation can arise by forging morphogenetic and metabolic partnerships that harness adjacent tissues to fuel newly emerged structures.

## Introduction

A well-established concept in biology is that evolution rarely invents new structures from scratch; instead, it frequently co-opts and reshufles existing underlying gene regulatory networks (GRNs) and developmental programs to generate novelty (1, 2). Classic examples of this principle include the evolution of beetle horns via the repurposing of appendage-patterning networks (3), the emergence of electric organs through the modification of existing myogenic programs (4), and the origin of mammary glands from ancestral cutaneous secretory units (5). In these cases, novelty arises primarily through the divergence or specialization of a single ancestral tissue type: patterning networks are redeployed to new locations, or existing cells (e.g., muscle fibers or skin glands) are modified to perform new functions.

However, the evolution of complex organs often requires a higher order of integration: the simultaneous recruitment and functional coupling of distinct tissue types that must cooperate to form a physiological unit. How does evolution orchestrate the assembly of such multi-component structures? Specifically, how are the developmental programs of disparate lineages—such as contractile muscle and secretory epithelium—”stitched together” to create a unified, functional novel organ? While the co-option of individual gene networks is well documented, the mechanisms by which evolution fuses these distinct modules into a metabolically and structurally cohesive “chimera” remain enigmatic.

The venom glands of spiders (araneae) provide an ideal model to address this challenge. A key innovation in the radiation of spiders, represented by over 50,000 species (6), these glands are not merely modified salivary tissues; they are high-pressure chemical weapons characterized by a unique architecture: a specialized secretory epithelium encapsulated in a muscular shell (7). This structure suggests a complex evolutionary origin involving the integration of distinct developmental inputs.

Here, we investigate the assembly of the venom gland in the spider *Parasteatoda tepidariorum*. We show that the gland is a developmental chimera, formed through the hierarchical recruitment of distinct ancestral networks. We demonstrate that the appendage-patterning gene *Distal-less* (*Dll*) is a key regulator of venom gland organogenesis, while the bHLH transcription factor Sage ensures secretory identity, repressing an ancestral neuronal fate. Crucially, we discovered that the myogenic transcription factor *sum-1* is not only involved in the development of the gland, but also drives the lipid metabolism and homeostasis of the secretory epithelium. Our findings suggest that the evolution of new complex organs proceeds not just by modifying single tissues, but by recruiting the metabolic and structural capacities of one tissue type to power the physiological demands of another.

## Results

### Transcription factors *Dll, sage* and *sum-1* are needed for normal venom gland development

Venom glands in spiders emerge within a *Dll* domain at the tip of the chelicerae and then migrate upwards to enter the cephalothorax during post-embryonic development. We previously found that the *Drosophila* salivary gland specific transcription factor *sage*, as well as the myogenic factor *sum-1* (ortholog of the *Drosophila nautilus* and vertebrate *myoD*), were expressed in adult venom glands as well as early in venom gland development (8). In spider embryos, *sage* is exclusively expressed in venom gland primordia, while *sum-1* is more broadly expressed in the chelicerae and other appendages (Fig.1A,B). Here, we performed parental RNA interference (pRNAi) to investigate the role of these three transcription factors in venom gland development (Fig. 1C). Phenotypes were assessed at three developmental stages – post-embryo early, first instar, and adulthood; here, we show results for first instars and adults, while results for post-embryo early are reported in SI Appendix.

**Fig. 1.**
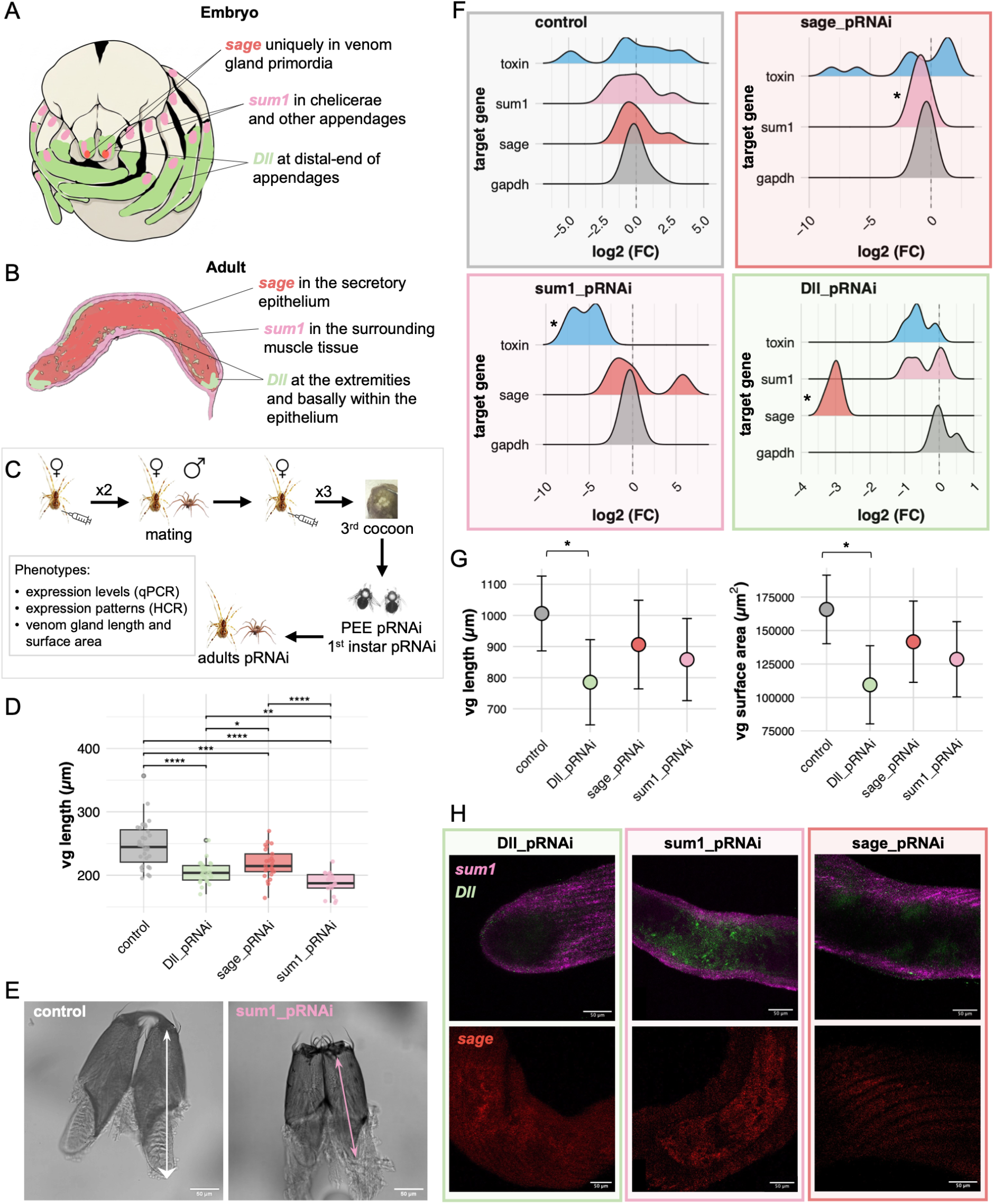
Effect of downregulation (pRNAi) of target genes on the development of spider venom glands. (A) Schematic view of the embryonic expression pattern of the target genes in the appendages. (B) Normal expression pattern of the target genes in adult venom glands. Experimental design detailing the dsRNA injections administered to females for pRNAi. (C) (D) Effect of pRNAi on venom gland (vg) length in first instar knockdowns compared to control. (E) Venom gland length in first instars, measured from the base of the fang to the proximal end of the gland. Representative images from control and *sum-1* pRNAi dissected chelicerae with glands. (F) Relative expression levels (log2 fold change) for the downregulated genes, the housekeeping gene *gapdh* (included as a control), and toxin gene *LOC107442349* in first instars. (G) Long-term effect of pRNAi in adult spiders measured as estimated marginal means (EMMs) of venom gland (vg) length and surface area. (H) Expression patterns of target genes in the venom glands of adult pRNAi spiders. All the results shown here are from individuals from the third laid cocoons following injection which had the highest phenotype observed in *Dll-*pRNAi where 100% of the spiderlings had 6 legs.

We discovered that in all knockdown first instar spiders, venom glands were significantly smaller than controls. *Sum-1* pRNAi had the strongest effect, with the least developed glands, followed by *Dll* pRNAi (Fig. 1D,E, SI Appendix, Table S1). Consistent with these phenotypic effects, qPCR showed that in *Dll* pRNAi, *sage* was strongly downregulated, while *sum-1* was weakly downregulated, with the latter statistically significant only in post-embryo early (Fig. 1F, SI Appendix, Fig. S1, Table S2). Similarly, after *sage* pRNAi, *sum-1* was downregulated, but *sage* expression was not significantly affected by *sum-1* pRNAi. While *Dll* and *sage* pRNAi had the most pronounced knockdown effect on developmental genes, *sum-1* pRNAi conversely produced the highest impact on the expression of the toxin gene *LOC107442349* (Fig. 1F). Because qPCR was performed on RNA extracted from whole individuals rather than isolated venom glands, the observed decrease in expression may reflect a reduction in overall gland size, and thus a lower proportion of gland-derived input RNA, rather than true transcriptional downregulation. We therefore corrected the expression values for the length of the venom glands, but the results did not change (SI Appendix, Table S2 and Fig. S2). Moreover, lower expression of target genes was also confirmed by RNA Hybridization Chain Reaction (HCR) (SI Appendix, Fig. S3). Thus all observations are consistent with downregulation of venom gland genes after target pRNAi. Similar results were observed also immediately after hatching in post-embryo early spiders (Table S2 and Fig. S1, SI Appendix).

### Gene knockdown via pRNAi has a long-lasting effect into adulthood

We allowed some knockdown spiderlings to grow and reach sexual maturity, and then we measured the total length and surface area of their venom glands. As in earlier stages, all the knockdown spiders had smaller venom glands compared to controls, with *Dll* pRNAi having the smallest glands, followed by *sum-1* and *sage* knockdowns, although only *Dll* pRNAi was statistically significant after false discovery rate (FDR) correction (Fig. 1G, SI Appendix, Table S3). Interestingly, pRNAi still affected gene expression, with low HCR signals for *sum-1* and particularly *sage* (Fig. 1H). Furthermore, *sum-1* pRNAi glands had abnormal *Dll* signals within the secretory epithelium, where they are typically absent in controls.

### *sage* knockdown results in downregulation of key venom biosynthesis components and upregulation of neuronal factors

Because *sage* plays a role in venom gland development, and its expression persists into adulthood where it is found exclusively in venom glands (8), we investigated its function in adult spiders. To this end, we silenced *sage* via RNAi and performed RNA-seq on dissected glands (Fig. 2A).

**Fig. 2.**
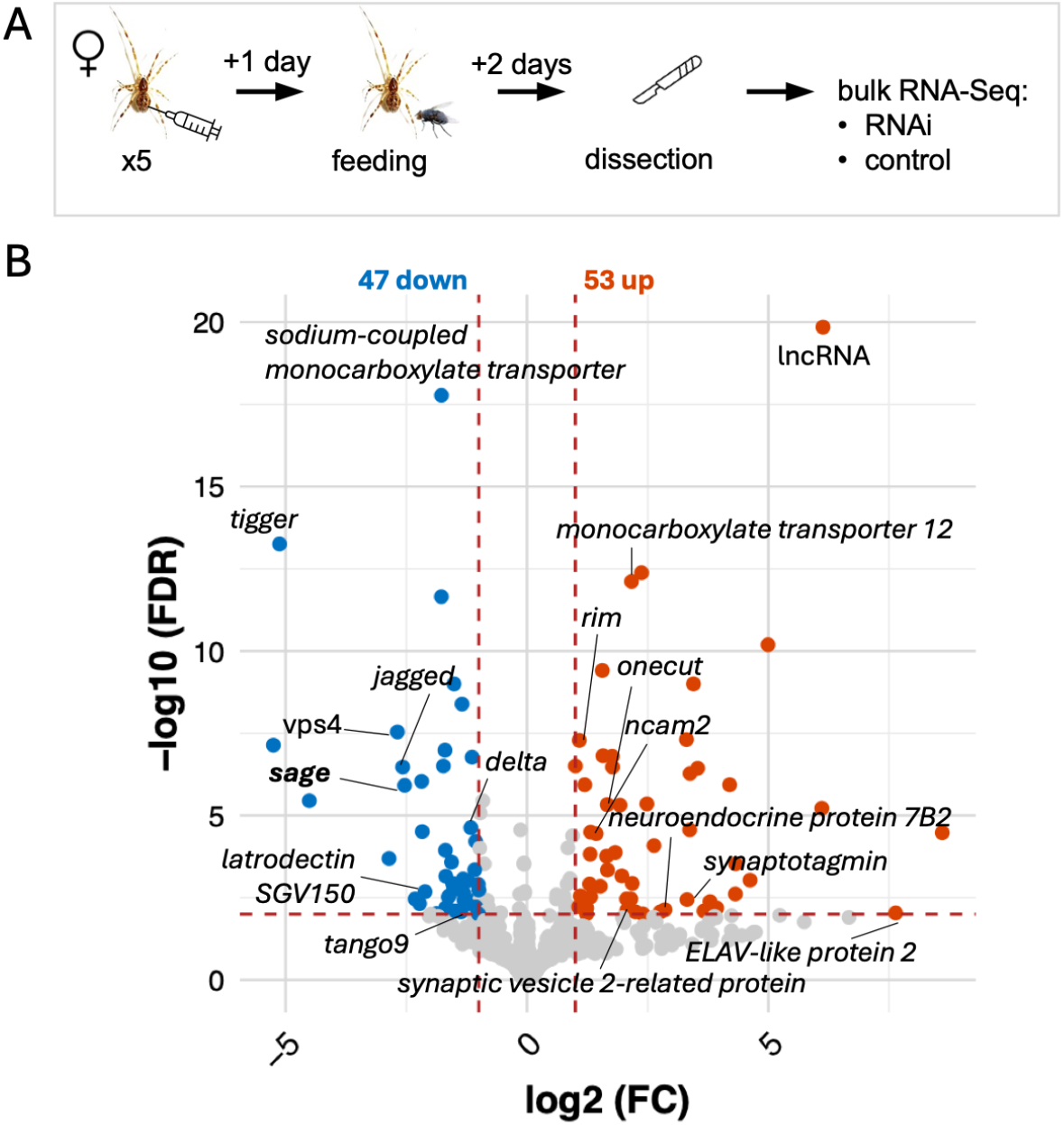
Effect of *sage* knockdown (RNAi) in adults on venom gland gene expression. (A) Overview of the experimental design. (B) Volcano plot of differential gene expression in venom glands between control and *sage* RNAi; differentially expressed genes (DEGs) with FDR < 0.01 and log_2_ fold change (FC) > 1 are highlighted in color, and the top DEGs are annotated (excluding genes named ‘uncharacterized proteins’).

A total of 47 genes were significantly downregulated, including *sage* itself, and 53 were upregulated (Fig.2B, SI Appendix, Fig. S4). Among the DEGs, there were only five toxin genes (four under- and one over-expressed), revealing that *sage* is not a major regulator of toxin expression. However, *sage* seems important for venom maturation and secretion. We found downregulation of chaperones and enzymes vital to toxin processing, such as latrodectins (*alpha-latrotoxin associated protein SGV150*), *neprilysin-2, cathepsin B-like*, and *hemocyte gamma-glutamyltransferase*. This was accompanied by downregulation of components of the secretory machinery, including metabolic transporterssuch as *sodium-coupled monocarboxylate transporter* and the *tigger transposable element-derived protein*, alongside core ER/Golgi trafficking proteins like *transport and Golgi organization 9* (*tango9*), *H(+)/Cl(-) exchange transporter*, and *vacuolar protein sorting-associated protein 4* (*vps4*). Finally, *sage* knockdown also caused disruption of cellular signaling with downregulation of the Notch ligands *jagged* and *delta*.

Conversely, genes upregulated upon *sage* knockdown revealed a shift toward a neuronal phenotype. These included key synaptic transmission components, such as *synaptotagmin 1, synaptic vesicle 2-related protein (sv2), regulating synaptic membrane exocytosis* (*rim*), and *neuroendocrine protein 7B2*, alongside cytoskeletal and adhesion regulators such as *rhophilin* (GTP-Rho-binding), *Rho GTPase-activating protein 18, liprin, WD repeat-containing protein 47, fasciclin 1* and 2 (orthologs to the vertebrates *neural cell adhesion molecule* (*ncam*)), and *ELAV-like protein 2* (*elavl2*). Finally, cell cycle control elements were also upregulated, notably apc, CDK inhibitors, ras association domain-containing protein, and Erk7/MAPK.

### Loss of *sum-1* causes downregulation of fatty acid anabolism and accumulation of lipid droplets in the secretory epithelium

The number of differentially expressed genes in *sum-1* knockdown glands was much greater than following *sage* RNAi, indicating that *sum-1* has a much broader regulatory network. Specifically, 463 genes were significantly downregulated, including *sum-1* itself, while only 79 upregulated, suggesting that *sum-1* acts as an activator rather than a repressor (Fig. 3A, SI Appendix, Fig. S5).

**Fig. 3.**
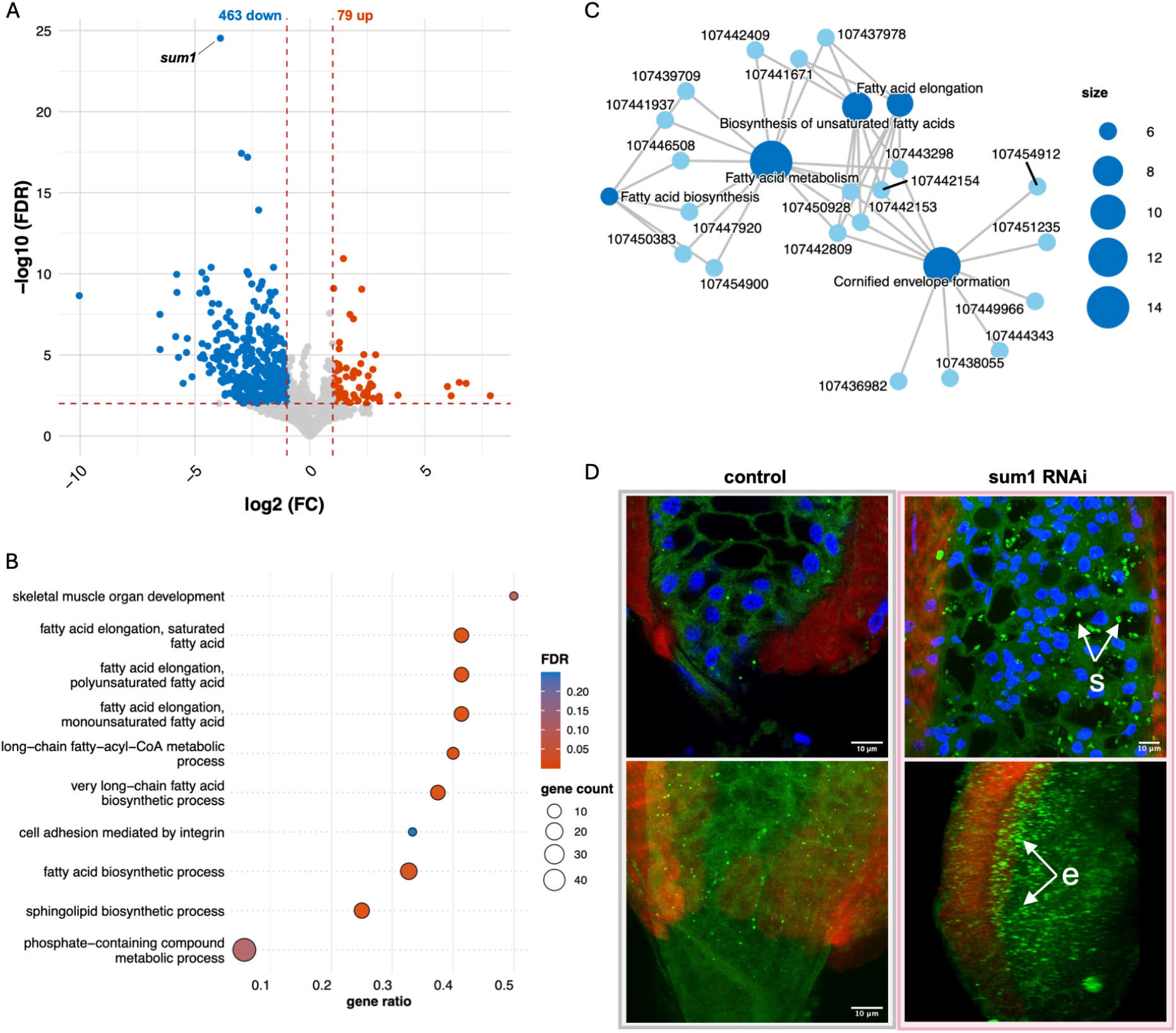
Changes in expression and lipid content of venom glands in *sum-1* knockdown (RNAi) adults. (A) Volcano plot of DEGs (FDR > 0.01, log_2_ fold change (FC) > 1) s in *sum-1* knockdown. (B) GO biological process terms enriched in downregulated genes. (C) Gene network of the top five enriched KEGG pathways. (D) Staining of neutral lipids (BODIPY, green), actin (Phalloidin, red), and nuclei (DAPI, blue) in venom glands of *sum-1* RNAi and control venom glands. Arrows highlight the accumulation of lipid droplets within secretory vesicles (s) and towards the basal membrane in the epithelium (e).

Functional enrichment analyses indicated strong inhibition of lipid biosynthesis and metabolic pathways (Fig. 3B,C). Downregulated genes were highly enriched for GO biological processes (BP) and KEGG pathways involving fatty acid metabolism, specifically “fatty acid elongation”, “long−chain fatty−acyl−CoA metabolic process”, “very long-chain fatty acid biosynthetic process”, “fatty acid and sphingolipid biosynthetic process”, “cornified envelope formation”, and “biosynthesis of unsaturated fatty acids”. Enriched molecular functions (MF)(“fatty acid elongase activity” and “fatty acid synthase activity”) and cellular compartments (CC) (“membrane”, “plasma membrane”, and “endoplasmic reticulum”) further supported this metabolic suppression. Individual downregulated targets included the muscle-associated genes *myosin-11* and *mef2C*, the homeobox genes *Mix*.*2* and *prospero*, and the specialized chaperone *Wnt ligand secretion mediator* (*Wntless*), which ranked among the top 20 mist significantly downregulated genes.

Among the 79 upregulated genes, we identified the homeobox *extradenticle* and genes associated with physiological stress. These included nearly all chains of the hemolymph oxygen-carrier *hemocyanin* (A, B, C, E, F), alongside enzymes driving metabolic mobilization: *arginine kinase, glycogen phosphorylase, phosphoglucose mutase 1*, and *ATP-dependent 6-phosphofructokinase*. Finally, a subset of genes were upregulated across both *sum-1* and *sage* RNAi treatments, including *ncam2*, a neuronal acetylcholine receptor, and the vesicle trafficking mediator *amphiphysin*.

Because of the strong effect of *sum-1* on fatty acids, we decided to stain lipid droplets in the glands. We used BODIPY, which is a fluorescence dye that stains neutral lipids within droplets, to test any potential differences between the knockdown and control groups (Fig. 3D). In normal venom glands, we found BODIPY stained small lipid droplets scattered exclusively across the secretory epithelium, with an even distribution. While most lipid droplet species where small dot-like structures, at times we observed larger, ring-like structures (SI Appendix, Fig. S6). In contrast, in *sum-1* knockdown venom glands, the lipid droplets where much more abundant, and they seemed to accumulate within the secretory vesicles. Importantly, when doing the 3D rendering, we observed stronger accumulation of signal towards the basal lamina rather than distributed across the secretory epithelium (Fig. 3D).

## Discussion

Spider venom glands are complex evolutionary innovations, requiring the integration of discrete secretory and muscular tissues into a unified organ. Our functional characterization of *Dll, sage*, and *sum-1* reveals that venom gland organogenesis does not rely on the deployment of isolated tissue modules. Instead, it is governed by a gene regulatory network that co-opts an appendicular patterning framework, maintains exocrine lineage identity via active neuronal suppression, and recruits a classic myogenic differentiation program to drive epithelial metabolic logistics.

### Evolutionary recruitment of the proximal-distal axis for gland organogenesis

Our functional analysis reveals that knockdown of the homeobox gene *Dll*, a highly conserved regulator of distal appendage structures across metazoans, disrupts both limb anatomy (9–11) and the formation of internal glandular structures, resulting in reduced venom glands. This dual effect matches the physical origin of the glands. Because the primordium arises at the tip of the embryonic chelicera and subsequently elongates proximally into the cephalothorax (8), its development depends on the proximal–distal (P-D) appendage patterning axis under the control of *Dll* (10). This mechanism contrasts with insect salivary glands, which instead develop along the anterior–posterior (A–P) body axis under the control of *Sex comb reduced, Extradenticle*, and *Homothorax* (12).

Furthermore, *Dll* knockdown causes a long-lasting downregulation of *sage* that persists into adulthood. This permanent loss shows that *Dll* does not just act as a temporary spatial cue during early development. Instead, *Dll* initiates a genetic program required to establish and maintain secretory cell identity throughout the life of the spider. Finally, our embryonic data show that *Dll* knockdown also reduces the expression of the myogenic factor *sum-1*. This reduction demonstrates that the early P–D positioning system acts upstream of both the epithelial and muscular lineages of the gland. We conclude that *Dll* functions as a master regulator for the entire venom apparatus. By controlling both *sage* and *sum-1, Dll* coordinates the development of the distinct tissue layers - the inner secretory epithelium and the outer muscle layer - needed to form a functional, contracting organ.

### The salivary gland-specific transcription factor *sage* maintains venom gland identity

Our findings show that the transcription factor *sage* is required for proper venom gland development as its loss through pRNAi results in smaller glands. This phenotype is similar to that observed in *sage* knockdowns in other arthropod glands. In *Drosophila, sage* mutants display severely underdeveloped embryonic salivary glands (13), and in the silkworm *Bombyx mori*, the silk glands, which are evolutionary modifications of salivary glands, are shorter and with lower cell numbers in sage RNAi larvae (14). These parallels indicate that *sage* has a deeply conserved role in building salivary gland structures across arthropods.

Beyond its developmental role, *sage* is required to maintain gland identity. Silencing *sage* in adults via RNAi results in downregulation of major components of the toxin processing (latrodectins) and secretory (*tango9* and *vps4*) pipelines. This secretory collapse mirrors the phenotype observed in *Drosophila* larvae, where *sage* mutations cause a decrease in the expression of the enzymes processing glue proteins, the primary secreted products of those glands, as well as the glue proteins themselves (15, 16). Conversely, in spiders, *sage* knockdown does not decrease the expression of the toxin genes themselves neither during development, as seen in first instars, nor in adult glands. This shows that, while *sage* controls the cellular machinery needed to mature and secrete venom, it does not directly control for toxin gene expression.

Another key finding is the downregulation of the Notch ligands *delta* and *jagged* coupled with the overexpression of typical neuronal markers (*elavl2, fasciclins*) (17, 18) alongside critical regulators of neuronal differentiation, cell adhesion and synaptic signaling (*rim, sv2, liprin*). In *Drosophila* salivary glands, the *jagged* ortholog *Serrate* drives epithelial differentiation and maintains the distinct morphological boundaries between secretory cells and the connecting duct (19). More broadly across arthropods, Notch signaling acts as the primary genetic gatekeeper of the ectoderm, actively repressing neural differentiation to allow specialized epithelial and secretory fates to emerge (20). The loss of *delta* and *jagged* in spider glands indicates that *sage* is required to maintain this vital signaling framework.

The immediate consequence of Notch signaling collapse is a phenotypic reversion toward a neuronal identity. The suppression of a neuronal program also mirrors the shared evolutionary history seen in early metazoans, where venom-secreting cells (cnidocytes) and neurons develop from common a pool of neuroglandular stem cell progenitors (21). We propose that *sage* acts as an evolutionary genetic switch that represses the ancestral neural fate to allow a new specialized secretory identity to emerge.

### The myogenic factor *sum-1* maintains glandular homeostasis

The secretory epithelium maintained by *sage* is surrounded by a muscle layer where the myogenic factor *sum-1* is expressed. Knockdown of *sum-1* via pRNAi results in the smallest glands among all transcription factors investigated, showing that *sum-1* plays a critical role in venom gland development. Furthermore, loss of sum-1 also results in downregulation of one of the main venom components, toxin *LOC107442349*, which is generally already actively transcribed in normal conditions (8). However, this as well as other toxin genes were are downregulated upon silencing of *sum-1* via RNAi in adults, suggesting that sum-1 does not control toxin genes directly, in agreement with the distinct spatial expression patterns of sum-1 and toxin genes. A lower expression in first instars might be the result of a delayed maturation of venom glands due to their smaller size, or a postponed toxin expression program until the glands reach an appropriate position within the cephalothorax.

Beyond development, *sum-1* exerts an important regulatory role in mature adult glands. Its silencing alters a large network of downstream genes including muscle-structural genes (myosin-11), signaling pathways (*wntless*), but most of all, genes involved in fatty acid anabolism, and lipid trafficking and metabolism.

The venom gland is a high-output organ with extreme metabolic and energetic demands required to translate, synthesize, and package complex toxins into secretory granules (22). To maintain this vesicular turnover, the glandular epithelium requires an influx of lipids to build phospholipid bilayers. Additionally, lipids and lipid-derivatives, such as acylpolyamines and sphingomyelinase targets, are active constituents of spider venoms that must be incorporated into the venom payload (23). Because the epithelium cannot synthesize this entire lipid budget *de novo*, these structural and cargo lipids must be imported from the hemolymph across the surrounding basal muscular layer, which acts as a mechanical and signaling gatekeeper (24). Under wild-type conditions, this high-throughput system processes and exports lipids apically so rapidly that large basal lipid pools do not accumulate. Considering all of these observations, we can formulate two hypothesis.

#### Hypothesis 1

##### cell-autonomous regulation of lipid anabolism

One hypothesis is that *sum-1* acts as a direct, cell-autonomous transcriptional activator of lipid anabolic pathways. Orthologs of *sum-1* have functional roles extending beyond myogenesis. For example, MyoD directly binds and regulates genes involved in fatty acid oxidation and oxidative metabolism (25). Under this paradigm, losing *sum-1* would directly suppress the transcription of fatty acid biosynthesis genes. This metabolic drop could explain the parallel downregulation of *wntless*, a transmembrane cargo receptor indispensable for the secretion of Wnt proteins (26). Wnt ligands require covalent modification with palmitoleic acid by the ER enzyme Porcupine before *wntless*-mediated transport. A depletion of cellular fatty acids would prevent Wnt lipidation, rendering the *wntless* chaperone system obsolete and leading to its transcriptional suppression.

However, our spatial data present two paradoxes that challenge this direct regulation model. First, *sum-1* expression is restricted to the outer muscular layer of the venom gland. If *sum-1* directly controls lipid biosynthesis, its knockdown should trigger a lipid deficit within the muscle fibers. Instead, BODIPY staining shows that the disruption of lipid homeostasis occurs in the adjacent secretory epithelium. Second, if the *sum-1* knockdown halted *de novo* fatty acid synthesis, the gland should experience a lipid deficit. Instead, we observe an accumulation of neutral lipid droplets packed inside the vesicle lumens.

Because a primary block in lipid synthesis cannot account for the accumulation of lipid-filled vesicles in the epithelium, these data argues against a cell-autonomous mechanism.

#### Hypothesis 2

##### non-cell autonomous paracrine signaling model

Alternatively, the downregulation of lipid pathways may represent an indirect physiological consequence rather than a direct transcriptional outcome of *sum-1* regulation. Under *sum-1* knockdown conditions, structural muscle-related genes (*myosin-11*) are downregulated, indicating that the integrity of the underlying muscle fibers may be compromised. We propose that this mechanical degradation disrupts vital paracrine crosstalk between the muscular gatekeeper and the adjacent glandular epithelium.

Concurrently, the downregulation of *wntless*, an essential cargo chaperone required for Wnt secretion, points to a collapse of localized Wnt signaling gradients. These gradients are critical for establishing apicobasal cellular polarity, inter-tissue crosstalking, and driving epithelial secretory programs (27, 28). Wnt ligands typically traffic across extracellular matrices and the intervening basal lamina to bind Frizzled receptors on neighboring epithelial targets, acting as paracrine cues that coordinate tissue homeostasis and directional patterning (29, 30). Without these instructive directional coordinates, the epithelial cells lack normal polarity cues and continue to import hemolymph-derived lipids, but fail to traffic or process them efficiently within the tissue. Consequently, the lipids stall at the entry point, generating the dense, localized band of BODIPY-positive puncta observed at the basal margin of the gland.

This localized traffic jam triggers a secondary, non-cell-autonomous transcriptomic collapse. As accumulating lipids physically congest the basal endoplasmic reticulum and Golgi networks, the epithelium silence endogenous fatty acid biosynthesis, elongation, and sphingolipid pathways to prevent lipotoxicity (cellular damage from excess lipid accumulation) (31). At the same time, the subset of lipids that successfully enter the ER and Golgi continue to be packaged into emerging venom vesicles, explaining the enlarged, intra-vesicular BODIPY signals observed experimentally. Concurrently, upregulation of enzymes driving metabolic mobilization and hemocyanin chains point confirms a state of overall cellular distress.

## Conclusions

The spider venom gland represents a complex evolutionary novelty that requires the precise coordination of distinct developmental programs. Rather than evolving entirely new genetic architectures, this specialized organ arose by co-opting and rewiring highly conserved bilaterian transcription factors. By analyzing the functional roles of *Dll, sage*, and *sum-1*, our study provides a blueprint for how evolution bridges divergent tissue modules - appendicular patterning, ectodermal cell-fate determination, and myogenesis - into a unified, high-output functional unit.

## Material and Methods

### Parental RNA interference (pRNAi) experiment

We investigated the roles of *Dll, sum-1*, and *sage* in the organogenesis of venom glands by means of pRNAi following a protocol adapted from Akiyama-Oda & Oda (32) which involves generation of dsRNA from PCR amplicons and multiple injections into females.

To generate dsRNA we extracted total RNA from whole embryos and dissected adult venom glands using the Direct-zol™ RNA MiniPrep kit (Zymo Research). 1 µg total RNA was converted to cDNA in a 20 µl reaction with SuperScript™ III Reverse Transcriptase (Invitrogen). PCR amplification was performed using 20-40 ng of cDNA per 50 µL reaction using DreamTaq PCR Master Mix (ThermoFisher). For *Dll*, we used the primers from Setton & Sharma (33). For *sage* and *sum-1* we designed the primers using the NCBI-Primers tool with a T7 linker sequence (CTAATACGACTCACTATAGGGAG) to both primers (forward and reverse). All primer sequences are available in SI Appendix, Table S4. The identity of the PCR amplicon were verified by Sanger sequencing. The *Dll* amplicon was cut out from agarose gels and purified with the MinElute^®^ gel extraction kit (Qiagen), while the other PCR products were directly purified with the MinElute^®^ PCR purification kit (Qiagen). 500 ng of purified PCR amplicon was used as template for synthesis using the MEGAscript T7™ transcription kit (Invitrogen) to obtain dsRNA. The final dsRNA concentration was 2 µg/ul eluted in nuclease-free water.

For each gene, three virgin females were anesthetized with CO^2^ and injected in the opisthosoma with 4 µg of dsRNA per injection. Injections were performed using a FemtoJet 4i microinjector (Eppendorf). The females were injected a total of 5 times at alternate day (Fig. 1C). After the 2^nd^ injection, they were mated with a male. Phenotypes were recorded for the 3^rd^ laid cocoon which was the one where nearly 100% of spiderlings had 6 legs in *Dll* knockdown.

For phenotype screening, we collected 20-30 individuals at post-embryo early stage, first instars and adults and fixed them in 4% PFA and stored in 100% methanol at −20°C until use. Approximately 10 samples were flash-frozen in liquid nitrogen and stored at −80 then used for RNA extraction and qPCR.

### Gene expression levels by qPCR

Total RNA was extracted from whole PEE and 1^st^ instar individuals and then approximately 20 ng of it was used directly as template for qPCR performed using the iTaq™ Universal SYBR^®^ green one-step kit (Biorad) to evaluate the expression levels of sage, sum-1 and toxin LOC107442349 (the latter only in first instars as in PEE the toxin gene is not expressed yet). PCR primers were designed with NCBI primers and are listed in SI Appendix, Table S5.

Samples were run in duplicates, the housekeeping gene Rpl27 was used for normalization and gapdh was also included to validate that the knockdown affect only the target genes and is not due to a global effect on RNA levels or normalization artifacts.

Relative gene expression was quantified using the comparative ΔΔCq method. Briefly, for each sample, Cq values of the target gene were normalized to the housekeeping gene Rpl27 to obtain ΔCq (ΔCq = Cq_target − Cq Rpl27). Relative expression changes between the knockdown and control samples were calculated as ΔΔCq (ΔΔCq = ΔCq_KD − ΔCq_control). Fold change was computed as 2^−ΔΔCq, and values were expressed as log_2_ fold change (log_2_FC = −ΔΔCq). For first instars, we also calculated the length-corrected log2 FC to account for differences in venom gland size. This is because the input RNA was from the whole body of the spiders, and we have shown that venom glands were smaller in knockdown compared to control, therefore it could be that the observed lower expression in knockdown was due to a smaller size of the glands, therefore less input RNA from the venom glands, compared to control. To correct for organ size, fold changes were adjusted for organ size by incorporating organ length as a proxy for tissue mass. Specifically, length-corrected fold change was calculated by multiplying the FC by the ratio of organ length in knockdown versus control (FC corrected = FC * (length in knockdown/length in control)).

### Measurements of venom gland length

All measurements were taken on images which were systematically taken with the same objective (10X) and no digital zoom, and they were processed with the software Fiji (ImageJ 1.54p). Measurements were taken twice independently and the mean between the two observations was calculated and used for the analysis.

#### First instars

Length in dissected chelicerae with the attached glands were taken by drawing a straight line from the proximal end of the gland to the base of the fang. This because often the distal end of the gland, where the venom gland duct start, it was not clearly visible, while the base of the fang and the venom gland tip at the proximal end were consistently visible.

Pairwise comparisons between the control and the knockdown were performed using the pariwise_t_test command in R v.4.4.3 (34) using the false discovery rate (FDR) p value adjustment.

#### Adults

To measure the venom glands in adult we used two measurements. First, we drew a segmented line all along the middle of the gland from the base of the duct in the distal end to the far end tip of the proximal end. This because the adult glands are not straight but more of a sausage and sometime is twisted so it is difficult to measure as a straight line. Secondly, we drew the contour around the gland and measure the surface area. Since we had both females and males, we compared the length and surface area controlling for the sex first by fitting a linear model (length/area ~ condition + sex) where condition is the knockdown treatment with ‘control’ as the reference group. We then calculated the estimated marginal means (EMMs) for the conditions using the *emmeans* function in R and we compared them by running the contrasts of each knockdown condition vs. the control. The p-values were adjusted using FDR.

### HCR of pRNAi post-embryo early and dissected venom glands

Before HCR experiments, fixed post-embryo early spiders and adult dissected venom glands were rehydrated with a series of graded methanol/0.1 %PBS-Tween-20. The head of post-embryo early was dissected in 0.1%PBS-Tween-20 and post-fixed in 4% PFA at room temperature for 20 min, then washed in PBS-T. After these steps, we followed the HCR protocol by Molecular Instruments for whole-mount samples.

### RNAi experiments in adult spiders

For RNAi experiments in adults, we injected 4 µg of dsRNA in females as we did for pRNAi

,for a total of four injections. We fed the spiders one day after the last injections and then performed the dissections the following day, i.e., day 2 post last injection and day 1 post-feeding (Fig. 2A). The control group was treated the same but injected only with water.

#### Bulk RNA-Seq

Total RNA was extracted from dissected venom glands of adult wild type and knockdown individuals as previously, and 25 ng of RNA was used to prepare libraries with the Watchmaker Library Prep kit (Watchmaker Genomics) using a unique dual indexing strategy and following the official protocol. Sequencing was performed on an Element Biosciences Aviti with a 2×75 cycles sequencing kit Couldbreak FS High Output run as single read. An average of 50M reads were generated per library with an amplicon size of 150 bp.

Reads were quality-filtered using Fastp v0.22.0 (35), pseudo-aligned to the transcriptome from the CAS_Ptep_4.0 (GCF_043381705.1) genome assembly using kallisto 0.48.0 (36), and gene abundances imported in R using the *tximport* package (37). Only genes with at least one read in at least three samples were kept for further analysis. Differential expression analysis of knockdown vs control was performed with DESeq2 1.46.0. Significant genes were with FDR < 0.01 and log fold change > 1.

KEGG enrichment of the up- and down-regulated genes was performed with clusterProfiler 4.14.6 (38) while GO enrichment was performed with topGO 2.58.0 (39).

#### Lipid staining

Fixed venom glands were incubated overnight 4 °C with BODIPY (Alexa Fluor 488) and Phalloidin (Alexa Fluor 594), then washed and incubated with DAPI before mounting. All images were acquired as z-stack with a Stellaris 5 White Light Laser (Leica Microsystems) inverted confocal microscope as described previously (8).

## Acknowledgements

This work was supported by a Swiss National Science Foundation COST grant IZCOZ0_205460 to GZ. We thanks the personnel of the Genomic Technologies Facility for the RNA-seq experiments, the Cellular Imaging Facility for the confocal microscopy, and Koen Oost for assistance with the BODIPY staining.

