## Supplementary material for "A developmental chimera: co-option of appendage, secretory and myogenic programs underlies spider venom gland evolution": SI Appendix

**Table S1. Statistics of pairwise comparison of venom gland length measurements in control and knockdown first instar spiders.**

| group1 | group2 | n1 | n2 | statistic | df | p-value | fdr |
| --- | --- | --- | --- | --- | --- | --- | --- |
| control | *Dll^pRNAi^* | 31 | 26 | 5.914 | 44.847 | 4.23E-07 | 1.27E-06 |
| control | *sage^pRNAi^* | 31 | 25 | 3.703 | 52.523 | 0.001 | 0.001 |
| control | *sum1^pRNAi^* | 31 | 19 | 7.926 | 46.519 | 3.52E-10 | 2.11E-09 |
| *Dll^pRNAi^* | *sage^pRNAi^* | 26 | 25 | -2.235 | 43.354 | 0.031 | 0.031 |
| *Dll^pRNAi^* | *sum1^pRNAi^* | 26 | 19 | 3.278 | 38.211 | 0.002 | 0.003 |
| *sage^pRNAi^* | *sum1^pRNAi^* | 25 | 19 | 4.835 | 41.980 | 1.82E-05 | 3.64E-05 |

fdr = p-value after false discovery rate correction.

**Table S2. Pairwise t-test statistics of gene expression levels for each knockdown condition versus control in post-embryo early (PEE) and first instar spiders.**

| stage | gene | condition | p-value | fdr |
| --- | --- | --- | --- | --- |
| PEE | ***gapdh*** | ***Dll^pRNAi^*** | 0.332 | 0.332 |
|  | ***gapdh*** | ***sage^pRNAi^*** | 0.151 | 0.227 |
|  | ***gapdh*** | ***sum1^pRNAi^*** | 0.056 | 0.168 |
|  | ***sage*** | ***Dll^pRNAi^*** | 0.027 | 0.055 |
|  | ***sage*** | ***sum1^pRNAi^*** | 0.155 | 0.155 |
|  | ***sum1*** | ***Dll^pRNAi^*** | 0.046 | 0.048 |
|  | ***sum1*** | ***sage^pRNAi^*** | 0.048 | 0.048 |
| first instar | ***gapdh*** | ***Dll^pRNAi^*** | 0.749 | 0.749 |
|  | ***gapdh*** | ***sage^pRNAi^*** | 0.149 | 0.341 |
|  | ***gapdh*** | ***sum1^pRNAi^*** | 0.227 | 0.341 |
|  | ***sage*** | ***Dll^pRNAi^*** | 0.000 | 0.001 |
|  | ***sage*** | ***sum1^pRNAi^*** | 0.823 | 0.823 |
|  | ***sum1*** | ***Dll^pRNAi^*** | 0.727 | 0.727 |
|  | ***sum1*** | ***sage^pRNAi^*** | 0.317 | 0.634 |
|  | ***toxin*** | ***Dll^pRNAi^*** | 0.716 | 0.716 |
|  | ***toxin*** | ***sage^pRNAi^*** | 0.505 | 0.716 |
|  | ***toxin*** | ***sum1^pRNAi^*** | 0.005 | 0.015 |
| first instar  length-corrected | ***gapdh*** | ***Dll^pRNAi^*** | 0.411 | 0.411 |
|  | ***gapdh*** | ***sage^pRNAi^*** | 0.022 | 0.034 |
|  | ***gapdh*** | ***sum1^pRNAi^*** | 0.014 | 0.034 |
|  | ***sage*** | ***Dll^pRNAi^*** | 0.000 | 0.001 |
|  | ***sage*** | ***sum1^pRNAi^*** | 0.987 | 0.987 |
|  | ***sum1*** | ***Dll^pRNAi^*** | 0.426 | 0.426 |
|  | ***sum1*** | ***sage^pRNAi^*** | 0.171 | 0.342 |
|  | ***toxin*** | ***Dll^pRNAi^*** | 0.540 | 0.540 |
|  | ***toxin*** | ***sage^pRNAi^*** | 0.395 | 0.540 |
|  | ***toxin*** | ***sum1^pRNAi^*** | 0.003 | 0.010 |

fdr = p-value after false discovery rate correction.

**Table S3. Comparisons of venom gland length and surface area in adult knockdown versus control.**

| measurement | condition | estimate | SE | df | t ratio | p-value |
| --- | --- | --- | --- | --- | --- | --- |
| length | ***Dll^pRNAi^*** | 220.795 | 90.547 | 51 | -2.438 | 0.055 |
|  | ***sage^pRNAi^*** | 99.872 | 92.782 | 51 | -1.076 | 0.287 |
|  | ***sum1^pRNAi^*** | 148.101 | 88.828 | 51 | -1.667 | 0.152 |
| area | ***Dll^pRNAi^*** | 56257.135 | 19287.147 | 51 | -2.917 | 0.016 |
|  | ***sage^pRNAi^*** | 24118.908 | 19763.256 | 51 | -1.220 | 0.228 |
|  | ***sum1^pRNAi^*** | 37210.928 | 18921.045 | 51 | -1.967 | 0.082 |

**Table S4. Primer sequences for dsRNA amplicon.**

| **locus** | **primer** | **sequence** | **length (bp)** | **reference** |
| --- | --- | --- | --- | --- |
| ***Dll*** | Dll_F | 5-ATGCCCAGGCTTACCCTATT-3 | 819 | (1) |
|  | Dll_R | 5-TGTCCCATGAGGAGATAGGC-3 |  |  |
| ***sage*** | sage_s_F | 5-CCGAGTCCATCAGTGGTTCG-3 | 481 | this study |
|  | sage_s_R | 5-TTCGCATGTATGACTGCCCA-3 |  |  |
|  | sage_F | 5-TATCCGAGAATGCCCCACAG-3 | 577 |  |
|  | sage_R | 5-CTTCGCATGTATGACTGCCC-3 |  |  |
| ***sum1*** | sum1_F | 5-AACAGACCTTGCCACCAACA-3 | 871 | this study |
|  | sum1_R | 5-TGAATCTCTGTGGCTCTGCG-3 |  |  |
|  | sum1_F2 | 5-CCCCCTCTCACTTCAACACC-3 | 745 |  |
|  | sum1_R2 | 5-GCCTCCTGCGTTCTCTCATT-3 |  |  |

**Table S5. Primer sequences for qPCR.**

| **locus** | **primer** | **sequence** | **length (bp)** |
| --- | --- | --- | --- |
| ***toxin*** | TX4_F | 5-AGTGCGCCCTGGTAAAACAC-3 | 152 |
| ***(LOC107442349)*** | TX4_R | 5-CATCCTGCATGCAATGAGTGGA-3 |  |
|  | TX11_F | 5-CAAAGAAGTGCGCCCTGGTAA-3 | 103 |
|  | TX11_R | 5-TCTAAACAAGCAAGCGTAGAACTGA-3 |  |
|  | TX2_F | 5-GCCCAGAAGCAGTTTGTGAAAA-3 | 119 |
|  | TX2_R | 5-GCGCACTTCTTTGTGGCAAT-3 |  |
| ***sage*** | PtSage_F | 5-ACCAACTACGAAACCTCCCG-3 | 109 |
|  | PtSage_R | 5-GGAGCAGGCTGGAGCATTTA-3 |  |
| ***sum1*** | PtSum1_F | 5-AGCTCTTCCGCTTCCACATC-3 | 114 |
|  | PtSum1_R | 5-TTGGGGCGACATAAAGGCAC-3 |  |
| ***RpL27*** | PtRpL27_F | 5-CAGCAAAGAGGCGAAAAGCA-3 | 71 |
|  | PtRpL27_R | 5-GCTCTTGCCAGTCTTGTACCT-3 |  |
| ***gapdh*** | PtGapdh_F | 5-TGAGGAAGGTGGAATGTTGGT-3 | 76 |
|  | PtGapdh_R | 5-TGGAATACTGCTGGGCTTCA-3 |  |

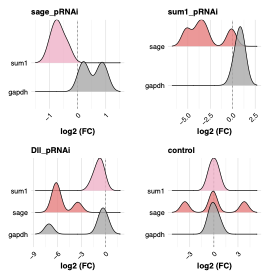

**Fig. S1. Relative expression levels (log2 fold change) in knockdown and control postembryo early spiders.** Expression levels were normalized to the housekeeping gene RPL27 and the fold change (FC) calculated using the ΔΔCq method. The housekeeping gene gapdh is included as a control and remains stable across conditions, supporting the specificity of the observed changes in target genes.

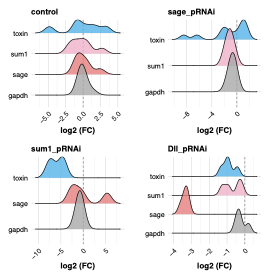

**Fig. S2. Relative expression levels (log_2_ fold change) in knockdown and control first instars adjusted for venom gland length.** Expression levels were normalized to the housekeeping gene RPL27 and the fold change (FC) calculated using the ΔΔCq method and corrected by the average length of venom glands in each condition. The housekeeping gene gapdh is included as a control and remains stable across conditions, supporting the specificity of the observed changes in target genes.

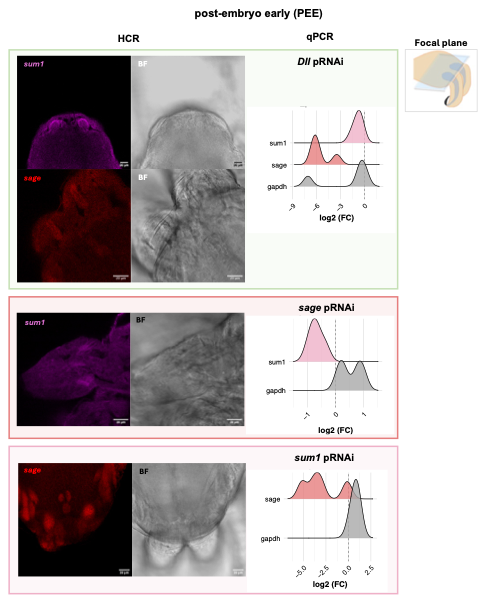

**Fig. S3. Hybridization Chain Reaction (HCR) signals of target genes in knockdown post-embryo early spiders.** On the left, HCR signals and brightfield (BF) images, on the right the corresponding relative expression levels (log_2_ fold change) from qPCR. A schematic of the focal plane is also displayed. Images from control individuals can be observed in (2).

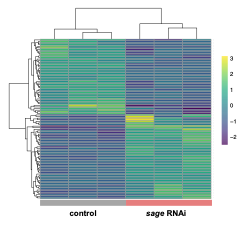

**Fig. S4. Heatmap of statistically significant differentially expressed genes (DEGs) in venom glands of *sage* knockdown (RNAi) vs control.** The heatmap displays the significantly upregulated and downregulated genes (fdr < 0.01, log2 FC > 1) (rows) between *sage* RNAi and control groups (columns). The values correspond to scaled and centered counts.

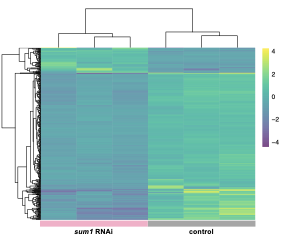

**Fig. S5.** **Heatmap of statistically significant differentially expressed genes (DEGs) in venom glands of *sum1* knockdown (RNAi) vs control.** The heatmap displays the significantly upregulated and downregulated genes (fdr < 0.01, log2 FC > 1) (rows) between *sum1* RNAi and control groups (columns). The values correspond to scaled and centered counts.

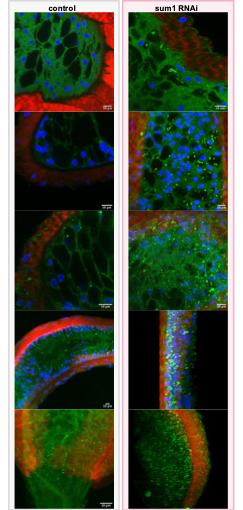

**Fig. S6. Lipid droplets staining in control and sum1 knockdown venom glands.** (A) Ring-like lipid structure in control venom gland. (B)
